# Assembly and annotation of two high-quality columbid reference genomes from sequencing of a *Columba livia* x *Columba guinea* F_1_ hybrid

**DOI:** 10.1101/2023.10.11.561892

**Authors:** Emily T. Maclary, Carson Holt, Gregory T. Concepcion, Ivan Sović, Anna I. Vickrey, Mark Yandell, Zev Kronenberg, Michael D. Shapiro

**Affiliations:** School of Biological Sciences, University of Utah, Salt Lake City, UT, USA; Department of Human Genetics, University of Utah, Salt Lake City, UT, USA; Pacific Biosciences, Menlo Park, CA, USA; Digital BioLogic d.o.o, Ivanić-Grad, Croatia

## Abstract

Pigeons and doves (family Columbidae) are one of the most diverse extant avian lineages, and many species have served as key models for evolutionary genomics, developmental biology, physiology, and behavioral studies. Building genomic resources for colubids is essential to further many of these studies. Here, we present high-quality genome assemblies and annotations for two columbid species, *Columba livia* and *C. guinea*. We simultaneously assembled *C. livia* and *C. guinea* genomes from long-read sequencing of a single F_1_ hybrid individual. The new *C. livia* genome assembly (Cliv_3) shows improved completeness and contiguity relative to Cliv_2.1, with an annotation incorporating long-read IsoSeq data for more accurate gene models. Intensive selective breeding of *C. livia* has given rise to hundreds of breeds with diverse morphological and behavioral characteristics, and Cliv_3 offers improved tools for mapping the genomic architecture of interesting traits. The *C. guinea* genome assembly is the first for this species and is a new resource for avian comparative genomics. Together, these assemblies and annotations provide improved resources for functional studies of columbids and avian comparative genomics in general.

**ARTICLE SUMMARY:** Pigeons and doves are important models for evolutionary genomics, developmental biology, physiology, and behavioral studies. Here, we present high-quality reference genome assemblies and annotations for two pigeon species, the domestic rock pigeon (*Columba livia*) and the African speckled pigeon (*C. guinea*). These assemblies and annotations provide improved resources for both comparative genomics and functional studies.

## INTRODUCTION

The domestic pigeon (*Columba livia*) is a unique model system for understanding the molecular and developmental basis of phenotypic diversity. *C. livia* has been under intensive artificial selection by pigeon fanciers for thousands of years, giving rise to more than 350 breeds with extensive variation in a variety of complex traits, including body size, beak morphology, feather color and morphology, and behavior (Stringham *et al*. 2012; Domyan and Shapiro 2017; Nimpf *et al*. 2019; Shao *et al*. 2019). In many cases, the magnitude of phenotypic diversity among *C. livia* approaches species-level diversity among wild bird species (Darwin 1868; Baptista *et al*. 1997). Thus, *C. livia* offers extraordinary phenotypic variation and experimental accessibility as a model for comparative genetics and developmental biology, both of which rely on a high-quality genome assembly and annotated gene models.

While domestic pigeons themselves are a spectacularly diverse single species, the family Columbidae is also one of the most diverse and geographically distributed extant avian lineages (Soares *et al*. 2016). Beyond *C. livia*, other doves and pigeons have served as important focal species for studies of physiology, behavior, and evolution (Johnson *et al*. 2010; Nimpf *et al*. 2019; Shao *et al*. 2019; Boyd *et al*. 2022; Burns-Cusato and Cusato 2022; Phillmore *et al*. 2022; Wasserman *et al*. 2023). Developing genomic resources for additional columbids provides resources for understanding avian diversity and ecological adaptation and critical tools for experimental biology. *Columba guinea,* the speckled pigeon, is of particular interest as a study species for comparative genomics because *C. guinea* and *C. livia* have partially overlapping geographical ranges and are interfertile. *C. livia* and *C. guinea* can hybridize to produce fertile male F_1_s, and potentially fully fertile backcrosses after only a few generations (Taibel 1949). We previously found that *C. guinea* contributed to the evolution of domestic and free-living populations of *C. livia* through the introgression of a potentially advantageous pigmentation trait (Vickrey *et al*. 2018).

Prior work developed reference genomes for *C. livia* (Shapiro *et al*. 2013; Damas *et al*. 2017; Holt *et al*. 2018). However, updated sequencing technology has paved the way for substantial improvements in contiguity and completeness, as seen in a recently released *C. livia* assembly (Wang *et al*. 2023). Additionally, new assembly methods can produce haplotype-resolved assemblies from heterozygous individuals (Nurk *et al*. 2020; Cheng *et al*. 2021). Here, we report two genome assemblies from a single *C. guinea* x *C. livia* F_1_ hybrid individual: the first assembly and annotation for *C. guinea* and an updated *C. livia* assembly with major improvements in contiguity, a new annotation incorporating full-length transcriptome evidence, and scaffolding based on extensive genetic linkage data.

## METHODS

### Animal husbandry

Pigeons were housed in accordance with protocols approved by the University of Utah Institutional Animal Care and Use Committee (protocols 19-02011 and 22-03002). Blood was collected from adult animals by brachial wing venipuncture.

### Genome sequencing and assembly

DNA was extracted from whole blood using the Qiagen DNEasy Blood and Tissue Kit (Qiagen, Valencia, CA) for parental F_0_ *C. guinea* (female) and *C. livia* (male) individuals. Samples were treated with RNAse during extraction. Isolated F_0_ DNA was submitted to the University of Utah High Throughput Genomics Shared Resource for library preparation using the Illumina Tru-Seq PCR-Free library kit. The resulting libraries were sequenced on the Illumina NovaSeq platform. Whole blood from one male F_1_ *C. livia* x *C. guinea* hybrid sample was submitted to the University of Delaware DNA Sequencing and Genotyping Center for DNA extraction, library preparation, and sequencing on the PacBio Sequel II system.

We used a standard short-read trio binning pipeline with hifiasm (Cheng *et al*. 2021). Parental *k*-mers were identified and counted with yak (v. r55; https://github.com/lh3/yak), (Cheng *et al*. 2021) using the settings -k 29 -b20 -t 12. The parental *k*-mer databases and PacBio hifi reads were then provided to hifiasm (v. 0.16.1-r375) (Cheng *et al*. 2021) for assembly of both genomes.

Assemblies were screened for contamination by genetic material from other organisms using FCS-GX (v. 0.3.0) (NCBI; https://github.com/ncbi/fcs/wiki/FCS-GX). We identified and removed one 7.5-kbp contig, h2tg000968l, in the *C. livia* assembly and one 7.5-kbp contig, h1tg000935l, in the *C. guinea* assembly predicted to be fungal in origin. Assemblies were screened for sequencing adaptor contamination using FCS-adaptor (https://github.com/ncbi/fcs); adaptors at the end of contigs were trimmed. One *C. livia* contig (h2tg000073l) with an internal adaptor was predicted to contain contiguous sequence based on both linkage data and contiguous sequence assembled across the same region in *C. guinea*. For this contig, we used NCBI BLASTN (v. 2.13.0) (Camacho *et al*. 2009) to identify reads from both the *C. guinea* and *C. livia* haplotype spanning the adaptor sequence. Adaptor sequence was excised and replaced with sequence from spanning reads with the *C. livia* haplotype. Other contigs with internal adaptors were split at the adaptor sequence. Based on these screens, contamination represented less than 0.001% of sequence in both genome assemblies.

We assessed the completeness of both assemblies with BUSCO (v. 5.3.2), using both the avian-specific aves_odb10 database and the more generalized vertebrate vertebrata_odb10 database (Manni *et al*. 2021).

### IsoSeq transcriptome sequencing

One *C. livia* embryo at the equivalent of Hamburger-Hamilton (Hamburger and Hamilton 1951) stage 25 was isolated from a fertilized egg and dissected in PBS. The head and body were separated and stored in RNAlater (Invitrogen, Cat. #AM7021). Total RNA was extracted from the whole head using the QIAGEN RNEasy Plus kit (QIAGEN, Cat. #74030). PacBio SMRTbell libraries were prepared, sequenced, and processed into hifi reads by the DNA Sequencing Center at Brigham Young University. Adapters were trimmed from hifi reads using lima (v2.5.0). Poly-A tails and artificial concatemers were removed using the isoseq3 (v3.4.0) “refine” command, and isoforms were clustered using isoseq3 cluster.

### Genome annotation

A repeat library generated for the previously published Cliv_2.1 annotation set (GCA_000337935.2) was used to generate a repeat annotation of both genomes, supplemented by RepBase RepeatMasker Edition version 20181026 (Bao *et al*. 2015; Holt *et al*. 2018) (http://www.repeatmasker.org).

MAKER (Holt and Yandell 2011) was used to annotate gene models for both *C. livia* and *C. guinea*. Protein evidence used by MAKER includes annotated zebra finch proteins downloaded from RefSeq (GCF_003957565.2), annotated chicken proteins downloaded from RefSeq (GCF_016699485.2), and all UniProtKB/Swiss-Prot proteins (Hillier *et al*. 2004; Boutet *et al*. 2007; Warren *et al*. 2010). IsoSeq sequences from *C. livia* were provided to MAKER as transcript evidence in FASTA format. Previously published Cliv_2.1 mRNA-seq data (Holt *et al*. 2018) were aligned using STAR (Dobin *et al*. 2013) against both the *C. livia* and *C. guinea* reference genomes, then assembled using StringTie (Pertea *et al*. 2015) and provided to MAKER as aligned GFF3 formatted transcript features. We then followed the published MAKER Alternate Protocol 1 with StringTie models being provided as both transcript evidence (est_gff) as well as gene prediction models (pred_gff) (Campbell *et al*. 2014).

*Ab initio* gene prediction through MAKER was configured to use Augustus (Stanke *et al*. 2006), trained according to the published MAKER Support Protocol 1 (Campbell *et al*. 2014) using UniProtKB/Swiss-Prot models first aligned by BLASTP (Camacho *et al*. 2009) and Exonerate (Slater and Birney 2005). These models were then converted by MAKER into gene models based on maximum open reading frame in the alignments. These protein alignment-based models were converted to GenBank format using MAKER’s zff2genbank.pl script. We then followed Augustus’s training documentation to produce final training models for *C. livia* and *C. guinea*. After the initial annotation, all Augustus *ab initio* models were processed through InterProScan (Jones *et al*. 2014) to rescue models that contain known protein domains but were initially rejected by MAKER in accordance with MAKER Basic Protocol 5 (Campbell *et al*. 2014) to produce the maximum annotation set.

The *C. livia* MAKER “max” models were compared to *Gallus gallus* BLASTX protein alignments using the BEDTools “intersect” command (Quinlan and Hall 2010). This process identified transcripts with exons overlapping two or more *G. gallus* protein models, which may represent false merges of two or more protein products in the final *C. livia* annotation set. These models were then manually curated in WebApollo (Lee *et al*. 2013). 742 MAKER “max” models were split into two or more models in the final annotation. All split models were again processed through InterProScan as above to identify known protein domains. The *C. guinea* MAKER models were not manually curated.

### Whole-genome comparisons

The Cliv_3 and Cgui_1 assemblies were aligned to each other using D-GENIES with Minimap (v. 2.24) to generate a genome-wide dotplot, PAF format alignments, and BAM format alignment files to calculate coverage (Cabanettes and Klopp 2018; Li 2018).The Cliv_3 assembly was also aligned to Cliv_2.1 scaffolds for comparison using the same approach. The Cliv_2.1 assembly has a small number of scaffolds that span multiple Cliv_3 contigs. To evaluate the utility of Cliv_2.1 reference genome to fill gaps in the Cliv_3 assembly, we used Quickmerge to merge Cliv_2.1 contigs or Cliv_2.1 scaffolds with the Cliv_3 assemblies (Chakraborty *et al*. 2016) and determine if the breaks between Cliv_3 contigs contained N tracts or contiguous sequence in Cliv_2.1 scaffolds.

Reciprocal alignments from Minimap were used to screen for putative large scale (>100 kbp) duplications, deletions, and inversions. Sequence regions with no alignments between species were identified as possible deletions in the species used as the “query” sequence. Tandem Repeats Finder (Benson 1999) was used to identify tandem repeats in non-aligning sequence. Repetitive intervals identified by Tandem Repeats Finder were merged with repetitive intervals annotated by RepeatMasker. The BEDTools “subtract” function (v. 2.28.0) (Quinlan and Hall 2010) was used to compare non-aligning regions of the Cliv_3 and Cgui_1 assemblies to the coordinates of repeat intervals to identify any large non-repetitive regions that did not align between species (Benson 1999). Putative duplications were identified using the BEDTools “genomecov” function to assess coverage in BAM format alignments. To identify possible inversions, PAF format alignments were filtered to exclude regions of low sequence identity and regions with multiple alignments, grouped by Cgui_1 contig, Cliv_3 contig, and relative orientation using the BEDTools “groupby” function, then manually examined for changes in relative orientation within pairs of aligned contigs.

### Comparison of Cgui_1 and Cliv_3 gene content

Cgui_1 and Cliv_3 gene models were compared using NCBI BLASTP to identify transcript pairs that were reciprocal best BLASTP hits (Camacho *et al*. 2009). For transcripts without reciprocal best BLASTP hits (“unmatched genes”), we used LiftOff (v. 1.6.3) to map GFF format annotations onto the reciprocal species genome (Shumate and Salzberg 2021). LiftOff mappings were compared to each annotation using the BEDTools “intersect” function to identify exon overlap (Quinlan and Hall 2010). For a small subset of representative gene arrays identified by LiftOff, NCBI BLASTN (Camacho *et al*. 2009) was used to compare sequence identity and exon boundaries for genes within the array.

### Comparison of repeat annotation

GFF format repeat annotations were collapsed into non-redundant, non-overlapping intervals using the BEDTools “cluster” and “groupby” functions (Quinlan and Hall 2010). Summed interval lengths were used to calculate total repeat content and cumulative distributions of repeat element size. Repeat type classifications from the GFF annotations were used for all analysis of repeat type distribution.

Terminal intervals were defined as the first and last 5 kbp of sequence from each contig. For comparisons of repeat distribution, 10000 random 5-kbp intervals were generated using the BEDTools “makewindows” function (Quinlan and Hall 2010). Intervals were not filtered to remove overlapping windows. Terminal and random intervals from each species were scanned for repeat content using BEDTools “intersect” to identify the number of basepairs covered by annotated repeats. Repeat composition of intervals was calculated based on the number of basepairs covered by each repeat type.

### Identification and evaluation of tandem 28-mer repeats

Tandem Repeats Finder (Benson 1999) was used to characterize a repetitive element identified in the Cliv_3 and Cgui_1 repeat annotations as “MuDR2-TC”. Examination of sequence within the repeats annotated as “MuDR2-TC” identified a tandem repeating 28-mer with the consensus sequence TGTCACAAACCCCATTGGACAGCGTGTG. Jellyfish (v. 2.3.0) (Marçais and Kingsford 2011) was used to count 28-mers in several avian genome assemblies (*Columbina picui,* GCA_013397635.1; *Patagioenas fasciata,* GCA_002029285.1; *Pterocles gutturalis,* GCA_000699245.1; *Streptopelia turtur,* GCA_901699155.2*; Tauraco erythrolophus,* GCA_000709365.1) and unassembled Illumina short-read genomic shotgun sequences (*Patagioenas oenops,* SRS1476223*; Columba palumbus,* SRS1416881; *Columba rupestris,* SRS346866; *Patagioenas speciosa,* SRS1476204; *Macropygia mackinlayi,* SRS1476214; *Streptopelia decaocto*, SRS1476207*; Streptopelia picturata* SRS1476196; *Zenaida asiatica,* SRS1476211) to screen for this 28-mer repeat in other columbid and outgroup genomes.

### Linkage map construction and anchoring to revised assembly

Genotyping and linkage map construction for five F_2_ intercrosses of *Columba livia* were carried out as previously described (Domyan *et al*. 2016; Boer *et al*. 2021; Maclary *et al*. 2021). Briefly, genotyping by sequencing (GBS) data were generated from founders and F_2_ populations. GBS reads were trimmed using CutAdapt (Martin 2011), then mapped to the Cliv_2.1 reference genome reads using Bowtie2 (Langmead and Salzberg 2012). Genotypes were called using Stacks2 (v. 2.5.3) by running “refmap.pl” (Catchen *et al*. 2013). We constructed genetic maps using R/qtl v1.46-2 (www.rqtl.org; (Broman *et al*. 2003).

To compare the linkage maps to the Cliv_3 assembly, each 90-bp locus containing a genetic marker was parsed from the Stacks2 output file “catalog.fa.gz” and queried to the Cliv_3 assembly using nucleotide-nucleotide BLAST (v2.6.0+) with the parameters -max_target_seqs 1 -max_hsps 1 (Camacho *et al*. 2009). Hits were filtered to retain alignments with an E-value < 4E-24, a threshold that permits multiple SNPs in each short sequence.

We used an Old German Owl x Racing Homer (OGO x Hom) linkage map as our primary reference for anchoring (Boer *et al*. 2021). 5978 markers from the OGO x Hom linkage map mapped to the Cliv_3 assembly with an E-value < 4E-24, and hits on 245 Cliv_3 contigs. We then identified markers mapping to 66 additional contigs in our 4 other crosses. We used linkage data from contigs shared between maps to determine placement for these 66 contigs within the OGO x Hom map. Markers within contigs were ordered based on basepair position, and the relative orientations of contigs with >3 markers spanning at least 30 kbp were determined based on calculated recombination frequencies in linkage maps.

### RagTag scaffolding

*C. livia* genome contigs were further scaffolded based on the *C. guinea* assembly using the RagTag v2.1.0 *scaffold* function (Alonge *et al*. 2019). The order and orientation of *C. livia* contigs within RagTag scaffolds were then compared to linkage map anchoring results. RagTag scaffolding provided ordering and orientation information for 31 additional contigs. Where possible, RagTag results were used to orient contigs with fewer than 3 GBS linkage markers or poorly spaced linkage markers. Contig orientations based on RagTag were also compared to oriented contigs from linkage map anchoring, and regions where RagTag and linkage map orientation disagree are noted in Supplementary Table 2.

### Comparison to chicken genome assembly

Cliv_3 proteins were compared to *Gallus gallus* GRCg7b reference proteins using NCBI BLASTP with the parameters -max_target_seqs 1 -max_hsps 1 (Camacho *et al*. 2009). Alignments were filtered to keep only pairs that were reciprocal best hits. Reciprocal best hits were then grouped by the linkage group of origin for the Cliv_3 protein and the chromosome of origin for the *Gallus gallus* protein, and the proportion of reciprocal best hit proteins shared between each Cliv_3 linkage group and *G. gallus* chromosome was calculated to identify correspondences. Gene order was not considered.

## RESULTS AND DISCUSSION

### Genome assemblies for *C. livia* and *C. guinea*

We generated 58,482,263,241 bp of circular consensus sequences from an F_1_ hybrid *C. livia* x *C. guinea* sample, with an average sequence length of 15.8 kbp, for an estimated coverage of 22.8x per haplotype. We additionally generated over 96 Gbp of *C. livia* short-read sequencing and 95 Gbp of *C. guinea* short-read sequencing from F_0_ individuals, for an approximate genome coverage of 75x per parental sample. We used hifiasm to assemble haplotype-resolved *C. livia* and *C. guinea* genomes from an F_1_ hybrid individual (Figure 1). The final *C. livia* assembly is 1,283,322,617 bp in length and consists of 1273 contigs, while the *C. guinea* assembly is 1,302,215,470 bp and has 1147 contigs (Table 1). Whole-genome alignment of the Cliv_3 and Cgui_1 assemblies show clear 1:1 alignments for the majority of both genomes (85% and 84% of the Cliv_3 and Cgui_1 assemblies, respectively; Supplementary Figure 1).

**Fig. 1.**
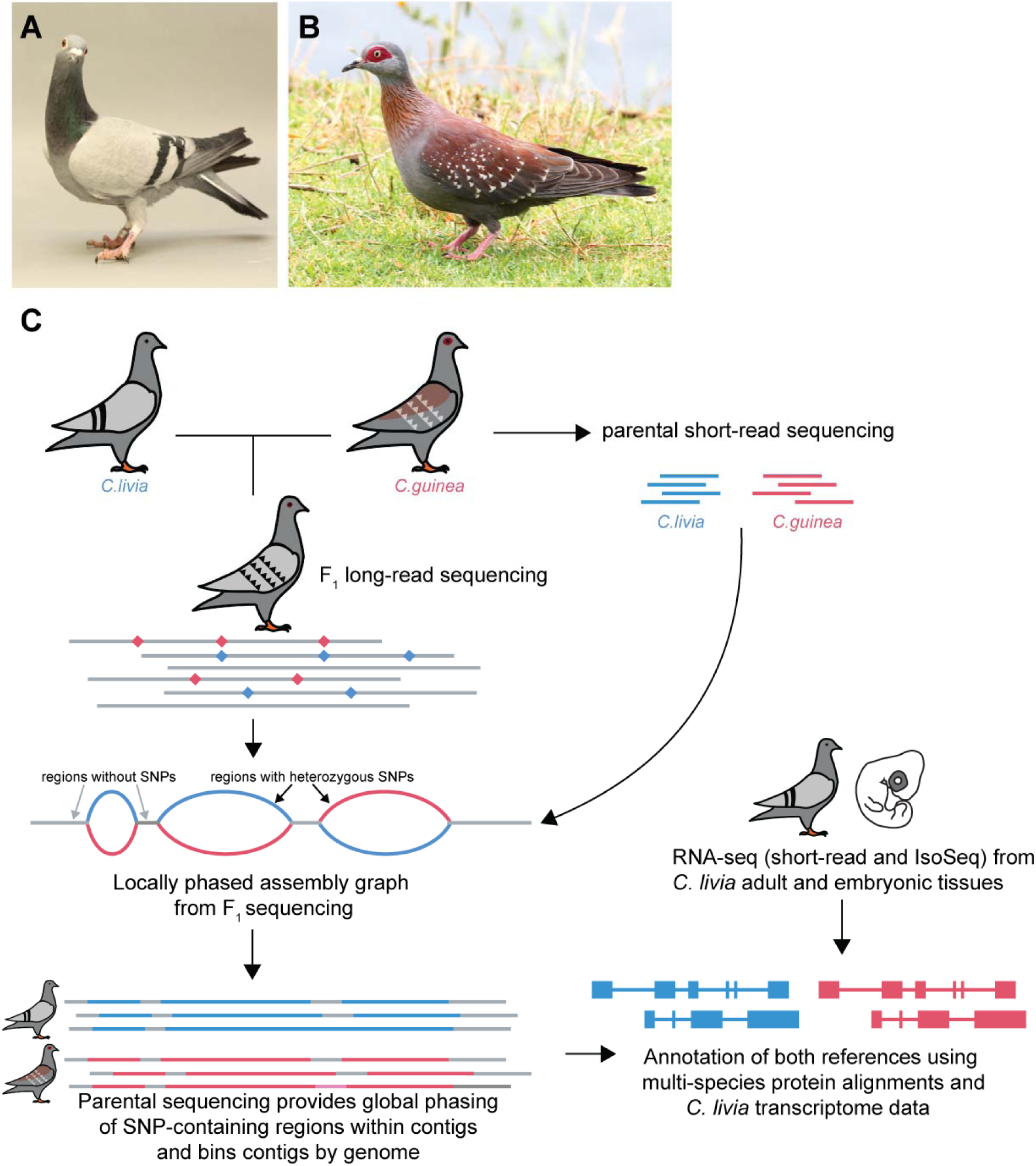
Genome assembly and annotation strategy. (A-B) Representative images of *Columba livia* (A) and *Columba guinea* (B). *C. livia* photo: Sydney Stringham. *C. guinea* photo: Roy Lowry, CC BY 4.0, via iNaturalist.co.uk. (C) Genome assembly strategy for haplotype-resolved *C. livia* and *C. guinea* genomes and annotations from an F_1_ hybrid.

**Table 1.** Genome and assembly statistics.

| Genome | Genome Size | Number of Contigs | Contig N50 | Annotated Genes | mRNA Models |
| --- | --- | --- | --- | --- | --- |
| Cliv_3 | 1.28 Gbp | 1273 | 6.97 Mbp | 16,853 | 18,431 |
| Cgui_1 | 1.30 Gbp | 1147 | 7.61 Mbp | 17,966 | 17,966 |
| Cliv_2.1<br>(Holt *et al.* 2018) | 1.11 Gbp | 107,177<br>(on 15,057 scaffolds) | 26 kbp | 15,392 | 18,966 |

The Cliv_3 and Cgui_1 assemblies have contig N50s of 6.97 Mbp and 7.61 Mbp, respectively. Therefore, the overall contiguity of the Cliv_3 assembly shows significant improvement over Cliv_2.1, which has a contig N50 of only 26 kbp (Table 1), and compares favorably to a recent *C. livia* PacBio-based assembly that has a slightly higher contig N50 of 7.8 Mb, but a larger number of contigs (1,949) (Wang *et al*. 2023). The Cliv_2.1 assembly contains several very large scaffolds, but they have numerous gaps, and the assembly is dominated by smaller scaffolds (Table 1, Figure 2). We examined Cliv_2.1 scaffolds that span multiple Cliv_3 contigs and found that gaps between Cliv_3 contigs that are spanned by Cliv_2.1 scaffolds always encompass one or more tracts of Ns (missing or ambiguous sequence). Thus, despite having several larger scaffolds, the Cliv_2.1 assembly does not contain contiguous sequence that can be used to resolve gaps between Cliv_3 contigs.

**Fig. 2.**
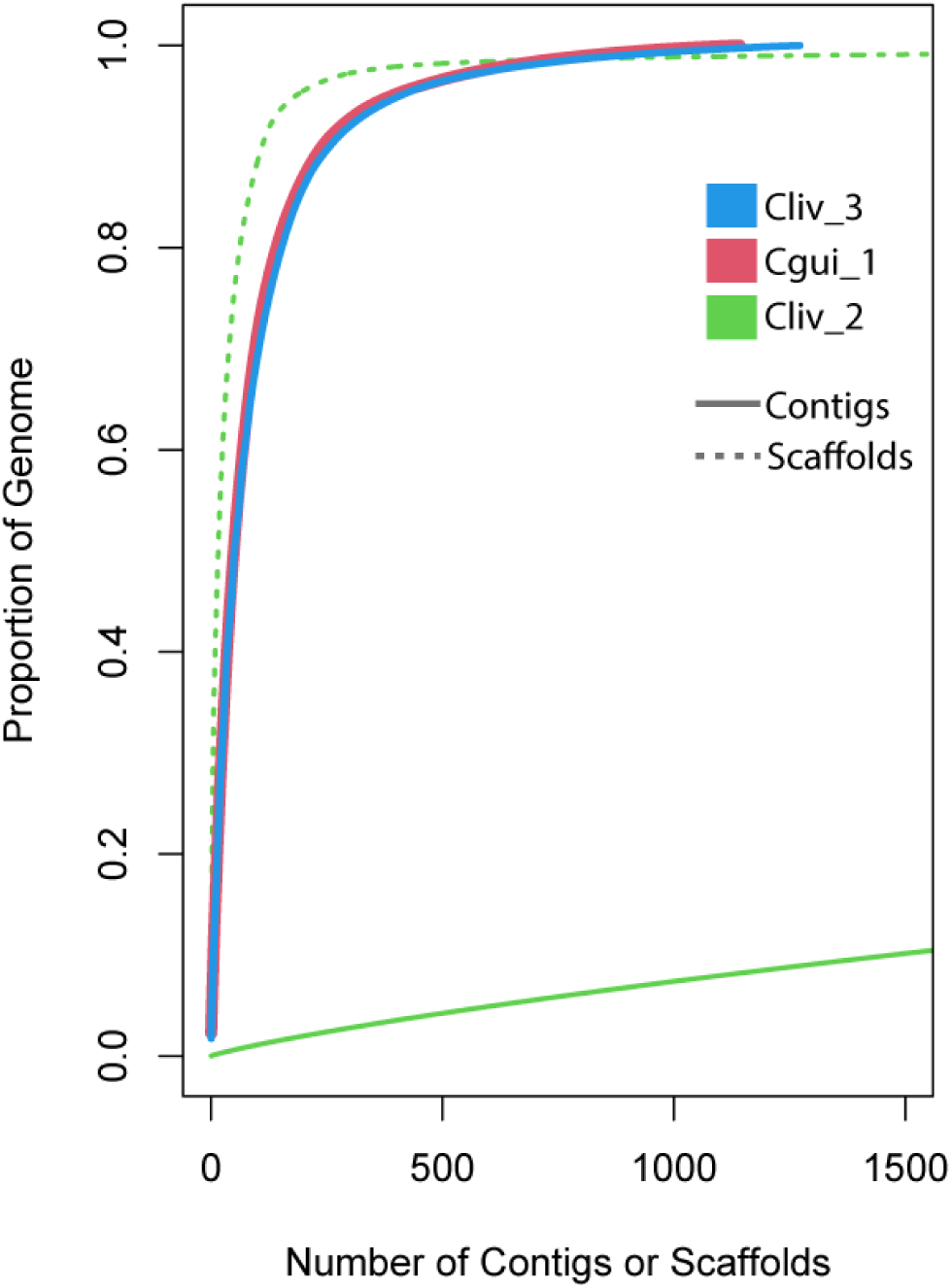
Cliv_3 and Cgui_1 assemblies show major improvements in contiguity over Cliv_2.1.

BUSCO analysis of highly conserved gene content of the new genome assemblies confirms both improved completeness and reduction in fragmentation of gene models compared to the Cliv_2.1 assembly. We found a 5% increase from Cliv_2.1 to Cliv_3 in the number of complete single-copy BUSCOs from the vertebrata_odb10 database, and a reduction in fragmented and missing BUSCOs (Table 2).

**Table 2.** BUSCO scores show improvements in completeness of Cliv_3 and Cgui_1 assemblies compared to the Cliv_2.1 assembly.

| Genome | BUSCO: Aves (aves_odb10 database) |  |  |  | BUSCO: Vertebrates (vertebrata_odb10 database) |  |  |  |
| --- | --- | --- | --- | --- | --- | --- | --- | --- |
|  | Complete (single copy) | Complete (duplicated) | Fragmented | Missing | Complete (single copy) | Complete (duplicated) | Fragmented | Missing |
| Cliv_3 | 97.0% | 0.4% | 0.4% | 2.2% | 95.6% | 0.7% | 1.6% | 2.0% |
| Cgui_1 | 96.9% | 0.4% | 0.5% | 2.2% | 95.5% | 0.8% | 1.6% | 2.1% |
| Cliv_2.1 (Holt *et al.* 2018) | 96.4% | 0.3% | 0.8% | 2.4% | 90.0% | 0.6% | 5.1% | 4.2% |

We examined whole-genome alignments of Cliv_3 and Cgui_1 for evidence of duplications, deletions, and inversions. We did not identify any contiguous >100-kbp regions with increased alignment coverage in either species, suggesting an absence of large duplications. We did identify several regions >100 kbp that did not align between species; however, >92% of this sequence content is repetitive. The lack of cross-species alignment in these regions could be due to their repetitive nature or a consequence of species-specific repeat expansion, and do not necessarily reflect large species-specific deletions or gains of unique sequences. Alignments between Cliv_3 and Cgui_1 show several large inversions (Supplementary Table 1), which could be characteristic of one or the other species, the Racing Homer *C. livia* breed, or the specific parents of the hybrid individual sequenced. Analysis of broader samples of both *C. guinea* and *C. livia* would be required to distinguish among these possibilities.

While contiguity is greatly improved in Cliv_3 and Cgui_1 relative to Cliv_2.1, the new genomes are not assembled to the chromosome level. We examined sequence at the termini (sequences at either end) of Cliv_3 and Cgui_1 contigs to determine if sequence characteristics contribute to assembly challenges and found an enrichment of annotated repeat elements compared to random 5-kbp intervals (two-tailed T test, p<1E-15). In addition to high repeat content, the terminal 5 kbp of Cliv_3 and Cgui_1 contigs showed changes in repeat composition compared to size-matched random windows across the genome, with enrichment of simple and satellite repeats (Supplementary Figure 2). Furthermore, only 0.27% of Cliv_3 repeat tracts genome-wide are >5 kbp in length. Contig termini represent less than 1% of the Cliv_3 genome, but 39% of these long repeat elements overlap the 5-kbp terminal intervals of contigs; 26% of terminal intervals show >99% coverage by a repeat tract longer than 5 kbp. Long simple repeats, satellites, and tandem repeat tracts can be particularly challenging for genome assembly, especially when repeat length exceeds read length or divergence between repeats is not sufficient to resolve them by phasing (Tørresen *et al*. 2019; Nurk *et al*. 2020; Peona *et al*. 2021). These 5-kbp intervals at the termini of contigs may represent fractions of larger repeat elements that we are unable to assemble because their lengths exceed common sequencing read lengths. We also noted that contigs from *C. livia* and *C. guinea* did not always terminate at orthologous sequences: in some instances, a single Cliv_3 contig spanned two Cgui_1 contigs, and vice versa, suggesting the presence of species-specific repeat expansions may have impacted genome assembly and contiguity.

### Linkage-based scaffolding of the Cliv_3 assembly

To evaluate the relative positions and orientations of Cliv_3 contigs, we used linkage data from F_2_ crosses and contigs from the Cgui_1 assembly that spanned gaps in the Cliv_3 assembly. We were able to assign 342 Cliv_3 contigs covering 1151 Mbp of sequence (89.7% of the Cliv_3 assembly) to 37 autosomal linkage groups and one Z-chromosome linkage group (Table 3, Supplementary Table 2). Several of the smaller (<20 Mbp) linkage groups in our anchored map consist of only 1 or 2 Cliv_3 contigs, suggesting that some contigs represent nearly complete chromosomes or chromosome arms.

**Table 3.** Linkage map anchoring of the Cliv_3 assembly identifies 29 autosomal linkage groups corresponding to 28 chicken chromosomes and one linkage group for the Z chromosome. *, chicken chromosome 4 is split into 2 autosomal linkage groups in pigeon, consistent with prior karyotyping and chromosome paint analysis.

| Linkage Group | Number of Contigs | Size (basepairs) | Corresponding *Gallus gallus* chromosome |
| --- | --- | --- | --- |
| 1 | 65 | 213,903,937 | 1 (196.45 Mb) |
| 2 | 59 | 163,618,457 | 2 (149.54 Mb) |
| 3 | 31 | 122,001,762 | 3 (110.64 Mb) |
| 4 | 12 | 79,237,614 | 4* (90.86 Mb) |
| 5 | 17 | 69,363,043 | 5 (59.51 Mb) |
| 6 | 6 | 55,545,771 | 7 (36.38 Mb) |
| 7 | 7 | 46,431,064 | 6 (36.22 Mb) |
| 8 | 5 | 35,246,159 | 8 (29.58 Mb) |
| 9 | 2 | 13,779,628 | 18 (11.62 Mb) |
| 10 | 2 | 15,514,691 | 20 (14.27 Mb) |
| 11 | 3 | 29,989,141 | 9 (23.72 Mb) |
| 12 | 4 | 20,545,355 | 13 (17.91 Mb) |
| 13 | 3 | 17,672,897 | 14 (15.33 Mb) |
| 14 | 2 | 21,279,300 | 11 (19.64 Mb) |
| 15 | 2 | 21,666,617 | 4* (90.86 Mb) |
| 16 | 6 | 7,135,027 | 24 (6.48 Mb) |
| 17 | 2 | 22,657,501 | 12 (20.12 Mb) |
| 18 | 2 | 11,857,108 | 17 (11.09 Mb) |
| 19 | 3 | 22,666,117 | 10 (20.45 Mb) |
| 20 | 3 | 14,961,191 | 15 (12.70 Mb) |
| 21 | 3 | 8,468,065 | 21 (6.97 Mb) |
| 22 | 1 | 8,321,153 | 26 (5.35 Mb) |
| 23 | 1 | 6,512,245 | 23 (6.25 Mb) |
| 24 | 4 | 6,093,282 | 22 (4.96 Mb) |
| 25 | 1 | 11,162,903 | 19 (10.46 Mb) |
| 26 | 3 | 6,907,984 | 27 (5.23 Mb) |
| 27 | 1 | 5,550,848 | 28 (5.44 Mb) |
| 28 | 3 | 3,442,929 | 25 (3.07 Mb) |
| 29 | 5 | 2,852,076 | 34 (3.47 MB) |
| Z | 70 | 84,521,162 | Z (86.04 Mb) |
| Other | 14 | 2,563,644 | NA |

Based on gene content, we identified Cliv_3 linkage groups that correspond to 29 chicken chromosomes (Table 3). Prior analyses showed that the ortholog of chicken chromosome 4 is split into two chromosomes in the rock pigeon (Derjusheva *et al*. 2004; Damas *et al*. 2017) and our linkage groups also support this split. We found at least one Cliv_3 linkage group corresponding to each chicken autosome >4 Mbp (n=26) and the Z chromosome. We were unable to identify a linkage group corresponding to several chicken microchromosomes <4 Mbp, perhaps due to genomic rearrangements between species.

### Genome annotations for Cliv_3 and Cgui_1

The Cliv_3 annotation consists of 16,853 gene models encoding 18,431 unique mRNAs (Table 1). The Cgui_1 annotation consists of 17,966 gene models and mRNAs (Table 1; only one isoform is annotated for each gene because we did not use species-specific transcriptome evidence). In addition to the short-read RNA-sequencing data used for the Cliv_2.1 annotation, the Cliv_3 annotation incorporates full-length IsoSeq transcript data as evidence. We assessed Annotation Edit Distance (AED), a measure of concordance between gene models and transcript evidence alignments, for the Cliv_3 and Cgui_1 datasets. AED scores were significantly lower for the Cliv_3 and Cgui_1 annotations than for Cliv_2.1, indicating improved concordance between MAKER annotation models and supporting *C. livia* RNA-seq evidence (Figure 3).

**Fig. 3.**
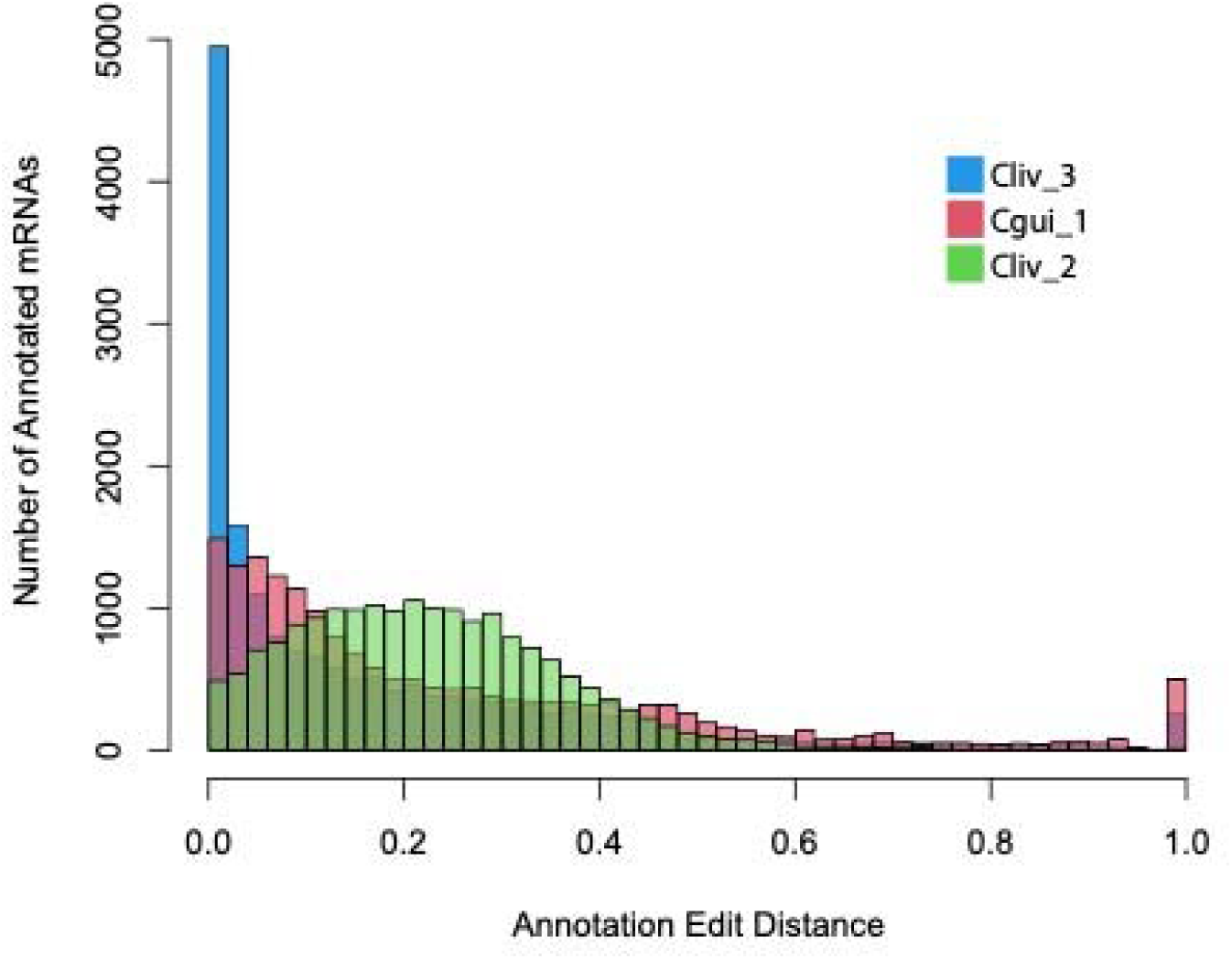
Annotation edit distances for Cliv_3, Cgui_1, and Cliv_2.1 assemblies. Cliv_3 and Cgui_1 show significantly lower AED scores (p < 1E^-^15, t-test) than Cliv_2, indicating improved concordance between annotation models and RNA-seq data in the new annotations.

We compared the MAKER *C. livia* and *C. guinea* annotation sets using reciprocal best BLASTP hit analysis and identified 13,821 matched gene models between the two annotations. Models not matched by reciprocal best BLASTP hits tend to have higher AED scores, indicating reduced concordance with protein and mRNA model evidence (Supplementary Fig. 3A). For the models not matched by best reciprocal BLASTP hits, we looked for correspondence between *C. livia* and *C. guinea* genes by using LiftOff (Shumate and Salzberg 2021) for reciprocal mapping between species. Ninety-four percent of non-matched Cliv_3 genes mapped to the Cgui_1 assembly, while 85% of non-matched Cgui_1 genes mapped to the Cliv_3 assembly. Most of these genes show shared intron-exon structure with a model from the alternate species assembly, but these correspondences were not identified by best BLASTP analysis for several reasons, including species-specific mutations that disrupted annotation. For example, 40% of Cgui_1 gene models that mapped to the Cliv_3 assembly did not have valid open reading frames, despite high sequence conservation between the two assemblies allowing for identification of orthologous exons. We also found evidence of fragmentation, in which a single gene model in one species is broken into several gene models in the other (Supplementary Fig. 3B). Interestingly, many of these non-matched and fragmented genes were also part of tandem arrays of genes with high sequence identity and conserved intron/exon structure (Supplementary Fig. 3B). Some of these non-matched gene models may also arise from species-specific expansion or contraction of tandem gene arrays.

### Differences in repeat content between *C. livia* and *C. guinea*

We found an expansion of annotated repeats in *C. guinea* compared to *C. livia.* In *C. guinea,* annotated repeats account for 20.8% of the genome assembly, while they account for only 16.8% in *C. livia* (see Supplementary Tables 3 and 4 for repeat annotations). The *C. guinea* genome is specifically enriched for longer repeat tracts (Fig. 4A). We also found changes in repeat composition (Fig. 4B). The Cliv_3 assembly is enriched for low complexity DNA, while the Cgui_1 assembly is enriched for simple and satellite repeats and DNA repeat elements. The higher DNA repeat content in Cgui_1 was particularly striking: this assembly contains nearly two-fold more sequence annotated as DNA repeat elements compared to Cliv_3 (16.15 Mbp in Cgui_1 vs. 8.23 Mbp in Cliv_3).

**Fig. 4.**
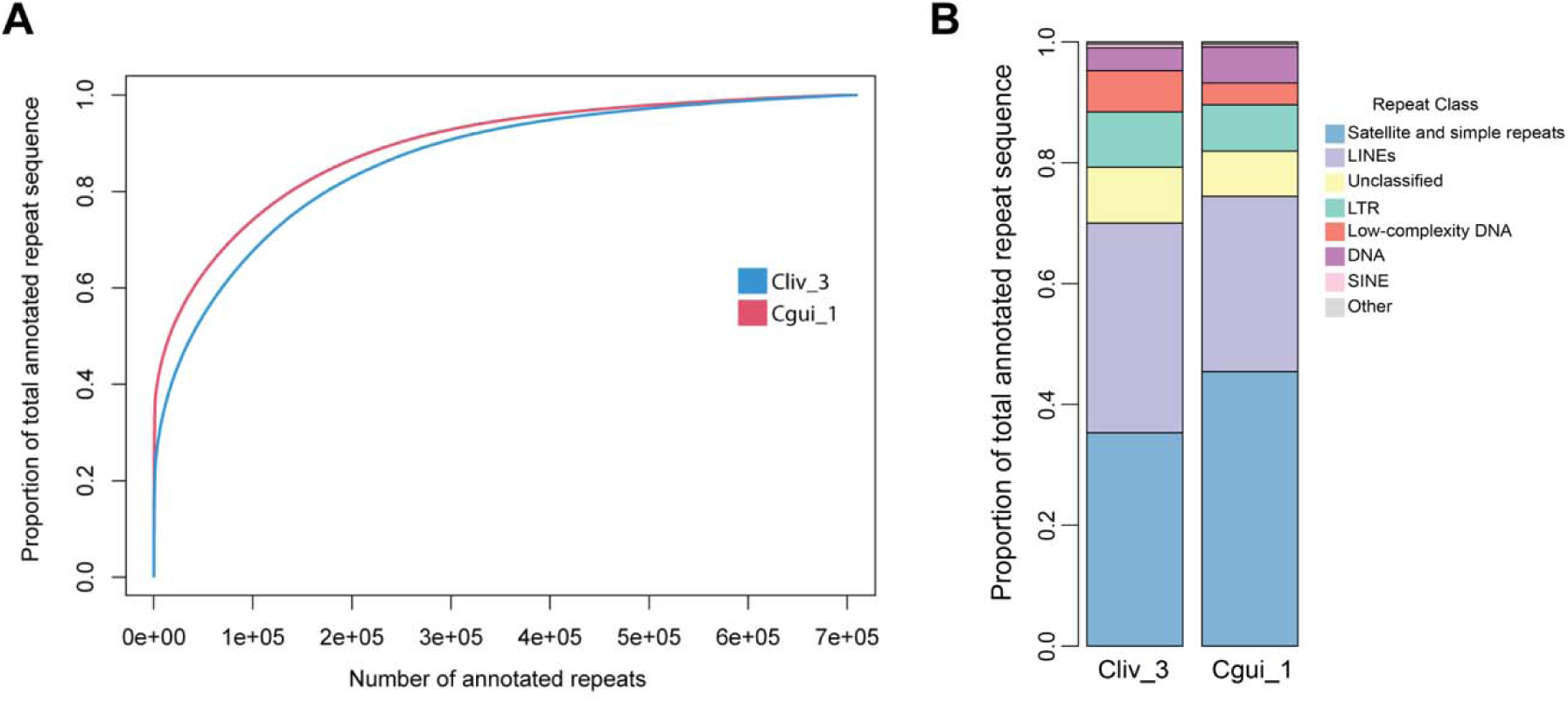
Repeat annotations differ between *C. livia* and *C. guinea.* (A) Cumulative distribution plot of *C. livia* and *C. guinea* repeats. The Cgui_1 genome is enriched for longer repeat tracts. (B) Distribution of repeat classifications in Cliv_3 and Cgui_1 assemblies.

The Cgui_1 repeat annotation showed substantial enrichment for a DNA element annotated as “MuDR2-TC”, which consists of tandem 28-mer repeats. BLAST hits to the NCBI nr database and screening for the 28-mer sequence in other avian genome assemblies and shotgun sequencing reads showed that this element is restricted to the monophyletic group that includes the columbid genera *Columba* and *Streptopelia* (Supplementary Fig. 4). The 28-mer repeats are frequently interspersed with PR1 elements that are found in *C. livia* centromeres (Solovei *et al*. 1996), and thus may represent centromeric sequence. However, the 28-mer repeat does not exclusively colocalize with PR1 sequences. Centromeric satellite sequence in columbid species is not well characterized. Unlike telomeric sequence, centromere satellite repeats are diverse and evolve rapidly between species, and changes in centromeric sequence may drive speciation (Fukagawa 2013; Melters *et al*. 2013). Whether this lineage-specific repetitive element in *Columba* and *Streptopelia* may play a role in the diversification of dove and pigeon species remains an open question.

### Advantages of parallel assembly and annotation of hybrid genomes

Annotation of a single assembly typically does not allow the types of comparisons that we are able to make between Cliv_3 and Cgui_1. Trio assemblies from other F_1_ hybrids either have not included annotations (Rice *et al*. 2019), or have focused analyses on specific genes of interest within the annotation (Low *et al*. 2020; Greenhalgh *et al*. 2022). As such, there is little precedent for comparing differences in annotations between closely related species with genome assemblies and annotations completed using the same pipeline. It is unclear how much variation in gene content should be expected between closely related species, particularly for tandem gene arrays or other repetitive elements, or how much cross-species comparisons could be affected by differences in assembly and annotation pipelines. While genome annotation pipelines and computational gene prediction perform well for conserved genes in non-repetitive regions that are well-supported by protein and RNA-seq evidence, the prevalence of non-matched gene models between the Cliv_3 and Cgui_1 assemblies, including fragmented models and arrays of gene models in repetitive regions, highlight both the continued need for manual curation and refinement of genome assemblies and the importance of focusing on well-conserved orthologs for assessment of genome completeness and interspecies comparisons.

## CONCLUSIONS

Here we present a new long-read genome assembly for *C. livia* and the first genome assembly for *C. guinea*, both derived from a single hybrid individual. The Cliv_3 assembly shows substantial increases in contiguity over the Cliv_2.1 assembly and resolves numerous gaps, and contiguity is similar in the Cgui_1 assembly. These increases in contig length and resolution of gaps will improve variant detection for comparative genomics in both species, including detection of structural variants or deletions that were previously hampered by short scaffold lengths or gaps that impair short-read mapping. The addition of the *C. guinea* genome will improve understanding of genomic diversity among columbids, a highly diverse and species-rich avian lineage. Furthermore, our updated Cliv_3 annotation shows improved concordance with RNA-seq and protein alignments, and the addition of long-read IsoSeq data provides new information about isoform diversity.

## DATA AVAILABILITY

This whole genome sequencing project has been deposited at DDBJ/ENA/GenBank under the accessions JAVLUT000000000 (*C. livia*) and JAVLUU000000000 (*C. guinea*). The versions described in this paper are version JAVLUT010000000 (*C. livia*) and JAVLUU010000000 (*C. guinea*). Sequencing data used to generate these assemblies are deposited in NCBI databases under BioProjects PRJNA1003477 (*C. livia*) and PRJNA1002478 (*C. guinea*). Short-read sequencing for *C. livia* is in the SRA database with sequence accession SRR25956418. Short-read sequencing for *C. guinea* is in the SRA database with sequence accession SRR25985494. PacBio sequencing of the F_1_ hybrid sample is in the SRA database with sequence accession SRR25956417. IsoSeq data generated for annotation is in the SRA database with sequence accession SRR26111786. Previously generated short-read RNA-seq data used for annotation is deposited in the SRA database with sequence accessions SRR5878849-SRR5878856. Repeat annotation GFF files for are available at FigShare as Supplemental Tables 3 (*C. livia* repeats) and 4 (*C. guinea* repeats).

## ACKNOWLEDGEMENTS

We thank Elena Boer for assistance with sample collection and processing. This work was supported by the National Institutes of Health (R35GM131787 to M.D.S.) and the H.A. & Edna Benning Foundation (M.Y.). The support and resources from the Center for High Performance Computing at the University of Utah are gratefully acknowledged.

